# Mechanistic pathway modeling reveals how IL-10 generates pleiotropic immune responses

**DOI:** 10.64898/2026.06.02.729479

**Authors:** Quim Martí-Baena, Carolina Segura-Morales, Jordi García-Ojalvo, Luis Serrano

## Abstract

IL-10 is a key anti-inflammatory cytokine whose activity is impaired in autoimmune diseases. However, IL-10 also promotes inflammation under certain conditions, limiting the efficacy of IL-10-based therapies. Because the principles underlying these opposing effects remain unclear, we developed a mathematical model of the IL-10 signaling pathway to understand how such pleiotropic responses arise. Considering that STAT3 signaling is buffered against changes in IL-10RB receptor affinity, we provide a minimal mechanistic explanation for IL-10 variants with anti-inflammatory or pro-inflammatory biased responses produced by an altered receptor affinity. By linking model-predicted pSTAT1 and pSTAT3 abundances with transcriptomic changes, we identified IL-10-responsive genes that are regulated at different pSTAT thresholds. This could explain how IL-10 elicits distinct downstream responses at different signaling strengths, and suggests that, depending on the required response, the affinity of IL10 for its receptors should not always be enhanced. Overall, our study highlights how modeling can help disentangle IL-10 pleiotropy to support the rational development of more effective IL-10-based therapies.

## Introduction

Cytokine-mediated communication is essential for coordinating immune responses across cell types (Saxton *et al*, 2021). Among these mediators, IL-10 plays a central role in limiting inflammation through diverse mechanisms such as suppressing the expression of pro-inflammatory cytokines or promoting the survival of FOXP3⁺ regulatory T cells (Wang *et al*, 2019b; Walter, 2014; Ouyang & O’Garra, 2019). Therefore, compromised IL-10 signaling is typically associated with inflammatory diseases such as inflammatory bowel disease (IBD)(Ouyang & O’Garra, 2019; Walter, 2014). IL-10-based therapies have thus been proposed to treat autoimmune diseases. These therapies, however, have shown no clear clinical benefits so far, as pro-inflammatory effects often limit their clinical translation (Tilg *et al*, 2002; Saraiva *et al*, 2020). Indeed, despite its canonical anti-inflammatory role, IL-10 can also stimulate monocytes and CD8⁺ T cells under specific conditions. So far, the only IL-10-based therapy that has advanced to a phase III clinical trial makes use of IL-10’s pro-inflammatory effects to target pancreatic cancer (Naing *et al*, 2018; Oft, 2014; Mühl, 2013; Wang *et al*, 2019b). Thus, although IL-10 is typically known for its immune inhibitory effects, IL-10 responses can be highly pleiotropic, with immune stimulation emerging depending on the target cell or IL-10 concentration (Naing *et al*, 2018; Wang *et al*, 2019b; Saraiva *et al*, 2020).

To understand the pleiotropic effects of IL-10, it is necessary to consider the molecular circuitry of IL-10 signaling. IL-10 is a dimeric cytokine. Therefore, its expressed monomeric form (IL-10M) is inactive due to improper folding (Montero-Blay *et al*, 2023; Walter, 2014; Saxton *et al*, 2021). To engage with its surface receptors, IL-10M must first dimerize (Figure 1A). After dimerization, each IL-10 subunit of the dimer independently binds one copy of the high IL-10 affinity, selectively expressed IL-10RA, and one copy of the low IL-10 affinity, ubiquitously expressed IL-10RB (Figure 1A)(Walter, 2014; Gorby *et al*, 2020). Although the affinity of IL-10RB for the IL-10 homodimer is very low (around 1 mM), it increases when the complex IL-10/IL-10RA is already formed, creating a two-step engagement with IL-10 (Josephson *et al*, 2001; Logsdon *et al*, 2002). The full IL-10 signaling complex (IL-10/IL-10RA/IL-10RB) allows the cross-phosphorylation of the receptor-bound kinases Jak1 and Tyk2, which subsequently phosphorylate specific tyrosine sites in the intracellular region of IL-10RA, where the downstream effector STAT3 can bind and be phosphorylated by Jak1 and Tyk2 (Figure 1A)(Riley *et al*, 1999).

**Figure 1:**
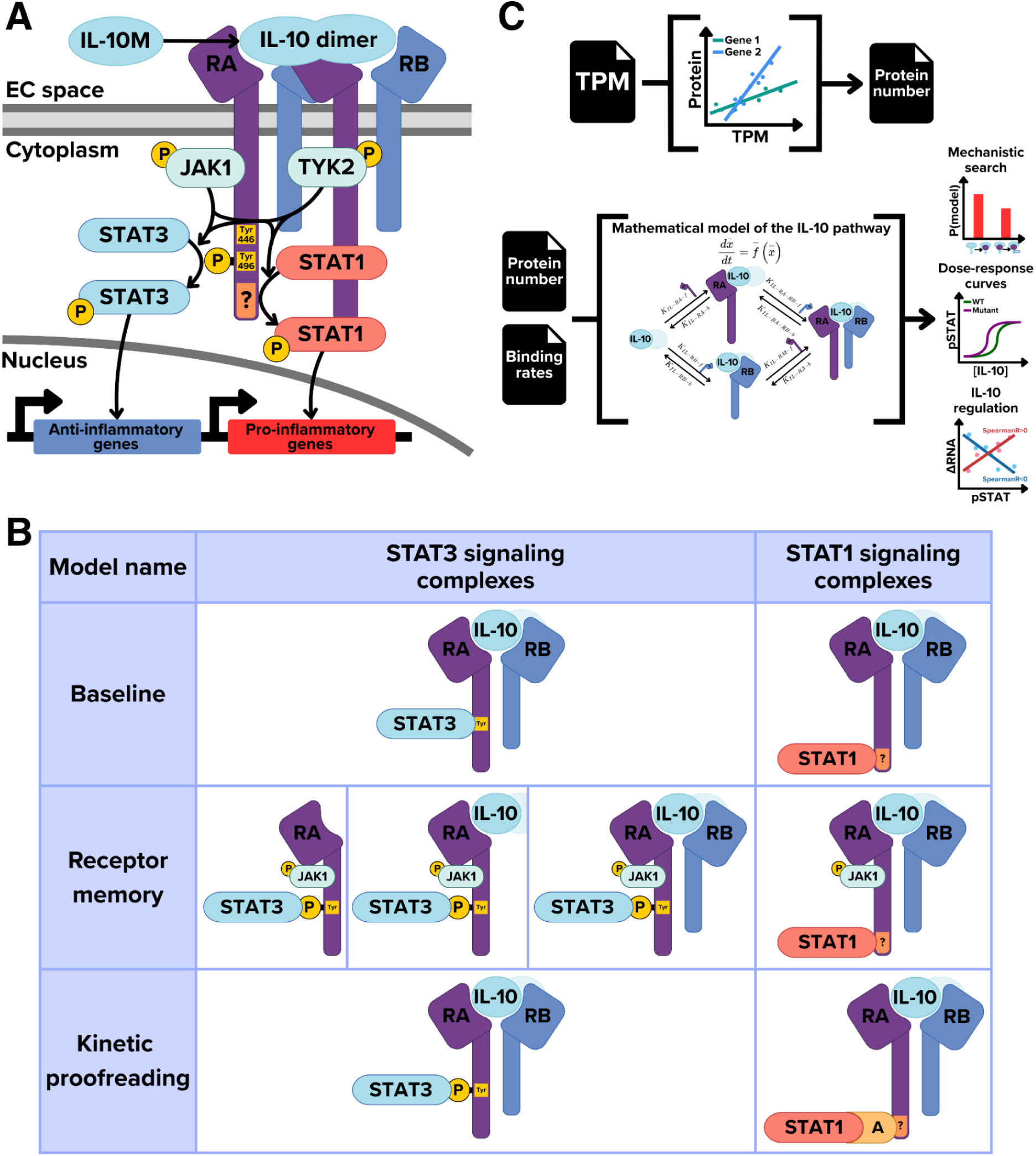
The IL-10 signaling pathway and modeling framework. **(A)** General view of the IL-10 signaling pathway. For IL-10 to bind to its surface receptors, first, it must form a homodimer. This allows the IL-10 dimers to bind to 4 surface receptors (2 IL-10RA and 2 IL-10RB), allowing the transphosphorylation of the receptor-associated JAK kinases. In return, these will phosphorylate tyrosine residues of IL-10RA, which enables the binding and phosphorylation of the transcription factor STAT3, which then migrates to the nucleus. Although STAT1 phosphorylation has been reported, its recruitment mechanism remains unresolved and is therefore represented schematically. **(B)** Table containing the protein complexes that can phosphorylate STAT3 or STAT1 across three simple mechanistic models to explore potential explanations for STAT1/STAT3 asymmetry. Only the binding of receptors to one subunit of the IL-10 homodimer is shown. For the Baseline model, STAT1 and STAT3 can be phosphorylated while the IL-10 signaling complex is formed (IL-10+IL-10RA+IL-10RB). For the Receptor Memory model, all complexes can signal through STAT3 as long as the IL-10RA-associated Jak1 remains phosphorylated. However, STAT1 still needs the IL-10 signaling complex at all times to be phosphorylated. For the Kinetic Proofreading model, STAT1 binds to IL-10RA via an adapter protein, whereas STAT3 binds IL-10RA directly. For a more detailed explanation of the models with a simplified reaction network, see Figure EV1. **(C)** Flow chart for our IL-10 signaling model. Transcriptomics data is used to estimate protein copy numbers via gene-specific linear models. These estimations are then used as an initial condition, in conjunction with our kinetic parameter set, to obtain the steady-state of phosphorylated STAT1 and STAT3 at different IL-10 doses using our IL-10 signaling model.

Although the mainly anti-inflammatory STAT3 is the central transcription factor mediating IL-10-induced downstream effects on target genes, IL-10-driven STAT1 activation has also been reported and is known to promote pro-inflammatory effects in monocytes (Rahimi *et al*, 2005; Weber-Nordt *et al*, 1996). While the precise mechanism by which STAT1 interacts with the IL-10 signaling complex remains unclear, prior studies place important constraints on this interaction. First, it has been shown that STAT1 does not interact via the phosphorylated tyrosine residues used by STAT3 (Weber-Nordt *et al*, 1996). Thus, STAT1 must interact with IL-10RA through a non-canonical site. This is consistent with prior work showing non-canonical interaction sites on the main receptor of other cytokines, including the C-terminal of IL-22RA1 with STAT3 and the acidic region of IL-2RB with STAT1 and STAT3 (Dumoutier *et al*, 2009; Delespine-Carmagnat *et al*, 2000). Finally, STAT1 signaling is more dependent on IL-10RB binding to IL-10 than STAT3 signaling, as IL-10 mutants with an increased binding affinity of IL-10RB to IL-10 increase STAT1 phosphorylation while maintaining WT IL-10 levels of phosphorylated STAT3 (pSTAT3)(Saxton *et al*, 2021; Gorby *et al*, 2020) (see also the Results section, Figures 3E-F).

The mechanism underlying the differential response of pSTAT1 and pSTAT3 to IL-10, however, remains unknown. Based on prior work in other cytokine systems, two candidate mechanisms may explain why STAT1 signaling depends more strongly on IL-10RB binding than STAT3 (Figure 1B, Figure EV1). The first mechanism follows a model introduced to disentangle different signals in IL-2 [19], where IL-10 receptors could retain IL-10-binding memory through the phosphorylation of their kinases (middle row in Figure 1B). Upon formation of the IL-10 signaling complex, the kinase associated with IL-10RA becomes phosphorylated, enabling STAT recruitment and activation (Walter, 2014). When either receptor dissociates, STAT binding and phosphorylation are assumed to cease as the receptor-associated kinases are dephosphorylated. However, dephosphorylation may not occur instantaneously. A dissociated IL-10RA could transiently retain a phosphorylated JAK1, thereby sustaining STAT3 phosphorylation independently of a fully assembled IL-10 signaling complex. In contrast, STAT1 signaling would remain dependent on the intact IL-10 signaling complex, due to a possible stronger requirement for the IL-10RB-associated Tyk2 kinase (Tyk2 Negatively Regulates Adaptive Th1 Immunity by Mediating IL-10 Signaling and Promoting IFN-γ-Dependent; Karaghiosoff *et al*, 2000). Another hypothesis (bottom row in Figure 1B) is that STAT1 binding would occur through an adapter protein (represented by A in the figure), as in a kinetic proofreading system (Hopfield, 1974). Since the interaction between STAT1 and IL-10RA would be weaker, high unbinding rates between IL-10 and IL-10RB would affect STAT1 more than STAT3, which instead would directly bind to the IL-10RA through its tyrosine sites. Some candidates for an adapter protein for STAT1 include PI3K, IFNGR1, or STAT3 (Herrero *et al*, 2003; Riley *et al*, 1999; Rahimi *et al*, 2005).

Mathematical modeling can help identify mechanisms that reproduce the differential STAT1/STAT3 behavior observed in experimental data. This has direct clinical implications, as optimizing IL-10 for anti-inflammatory activity likely requires favoring STAT3 signaling while limiting STAT1-driven responses. Using mathematical models of the IL-10 pathway and Bayesian inference, we aim to provide mechanistic reasoning for how opposing responses can occur in IL-10, focusing on dissecting the IL-10 response at two levels: pSTAT phosphorylation and downstream transcriptional programs. To do that, we developed an ordinary differential equation (ODE) model representing interactions in the IL-10 pathway, from IL-10 dimerization and receptor binding to different STAT1 and STAT3 binding and phosphorylation mechanisms (Figure 1C). By combining our mathematical model of the IL-10 pathway with cell-type-specific expression data of IL-10 receptors and STATs and kinetic parameters around experimental values, we reproduced the differential behavior of pSTAT1 and pSTAT3 observed in IL-10 dose-response curves. This methodology also allowed us to gain insights into the transcriptional responses downstream of the IL-10 pathway, revealing how IL-10 variants with an enhanced response may not confer additional effects over wild-type IL-10 for certain target genes. Overall, our study shows how computational modeling can shed light on the complexity of IL-10 responses and support the clinical development of IL-10-based therapies.

## Results

### Mechanistic models of IL-10 signaling

To simulate IL-10 responses, we developed a kinetic model that simulates the IL-10 pathway from IL-10 dimerization and receptor binding to STAT phosphorylation. This allows us to predict the steady-state pSTAT1 and pSTAT3 in a single cell from the cell-specific expression of receptors and STATs, the IL-10 variant-specific binding parameter values, and the extracellular IL-10 concentration. The reaction networks of the three proposed IL-10 signaling models, parameter meanings, and parameter values used in our IL-10 signaling model can be found in Figure EV1 and Appendix Tables S3 and S4, respectively. The ordinary differential equations (ODEs) of the model are included in the Appendix.

### Linear models infer protein copy numbers from RNA sequencing data

Differential cytokine responses are thought to arise from cell-type-specific expression of receptors and STAT molecules (Cui *et al*, 2024; Vento-Tormo *et al*, 2018; Ramilowski *et al*, 2015). Accordingly, to simulate responses in a given cell type, our mechanistic models require estimates of the copy numbers of IL-10 pathway proteins per cell, which are rarely available and limit applicability. To accomplish this, we built four gene-specific linear models (IL-10RA, IL-10RB, STAT1, STAT3) to infer protein copy number from mRNA levels. To train these models, we combined published transcriptomic and proteomic measurements from immune cell types (Sikkema *et al*; Rieckmann *et al*, 2017; von Haehling *et al*, 2015; Aizaki *et al*, 2021; Jurlander *et al*, 1997; Tian *et al*, 2014; Sikkema *et al*, 2023) with publicly available transcriptomic data across cancer cell lines (Barretina *et al*, 2019; Data ref: Barretina *et al*, 2019). Given the limited number of cell types with measured IL-10 receptor abundances, we also quantified IL-10RA and IL-10RB protein copy numbers across four cancer cell lines using spike-ins of heavy peptides (see Methods for absolute proteomics data generation).

These models yielded a low mean absolute error (*MAE*) in the training dataset across all 4 genes (Figure 2A), but their performance decreased on an independent dataset of 29 healthy human tissues used as validation (Appendix Figure S1)(Eraslan *et al*, 2019). However, the higher error in the prediction of protein counts in unseen conditions of the linear models did not significantly affect the performance of our downstream IL-10 signaling model (described below), when comparing cell types where the receptor/STAT copies per cell were quantified experimentally versus the case in which the copy numbers were estimated with our linear models (Appendix Figures S2A-B). These results support the use of transcriptomic-based estimates to enable a broadly applicable prediction of cell-type-specific responses with mechanistic models.

**Figure 2:**
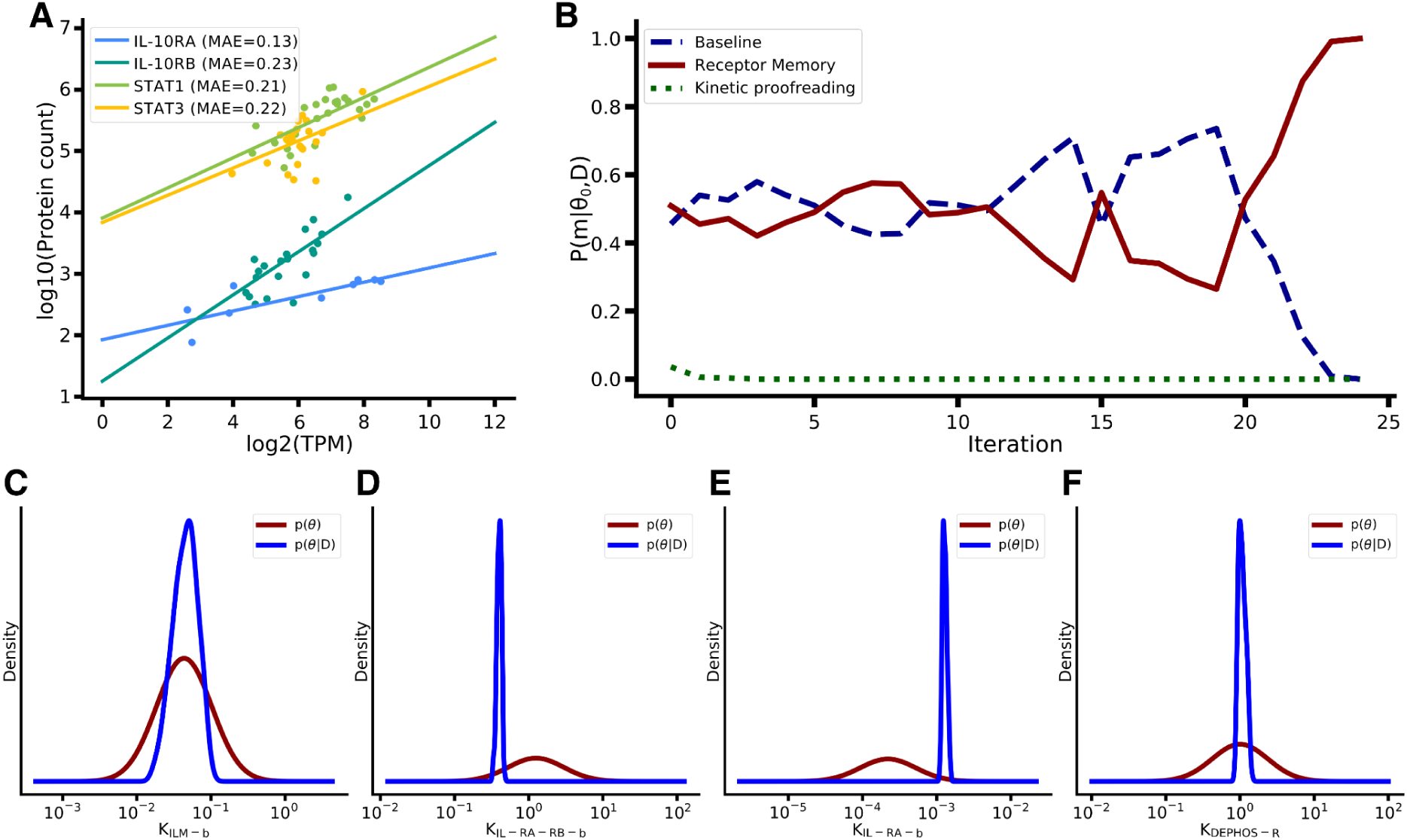
Fitting of the RNA to Protein linear models and model selection with parameter search via ABC-SMC. **(A)** Gene-specific linear models predict the number of proteins per cell of a gene with the level of messenger RNA molecules of the same gene. RNA abundance is measured in transcripts per million (TPM) whereas protein abundance is measured in protein counts per cell. **(B)** Probability (*p*(*m*|θ_0_, *D*)) of each of the 3 proposed models across ABC-SMC iterations. **(C, D, E, F)** ABC-SMC-derived prior and posterior distributions of the parameters (C) IL-10 dimer dissociation rate (*K*_*ILM*−*b*_), (D) unbinding rate of IL-10RB to the IL-10+IL-10RA complex (*K*_*IL*−*RA*−*RB*−*b*_), (E) unbinding rate of IL-10RA to IL-10 (*K*_*IL*−*RA*−*b*_) and (F) IL-10RA dephosphorylation rate (*K*_*DEPHOS*−*R*_) for WT IL-10. All reaction rates are displayed in 1/*s* units

**Figure 3:**
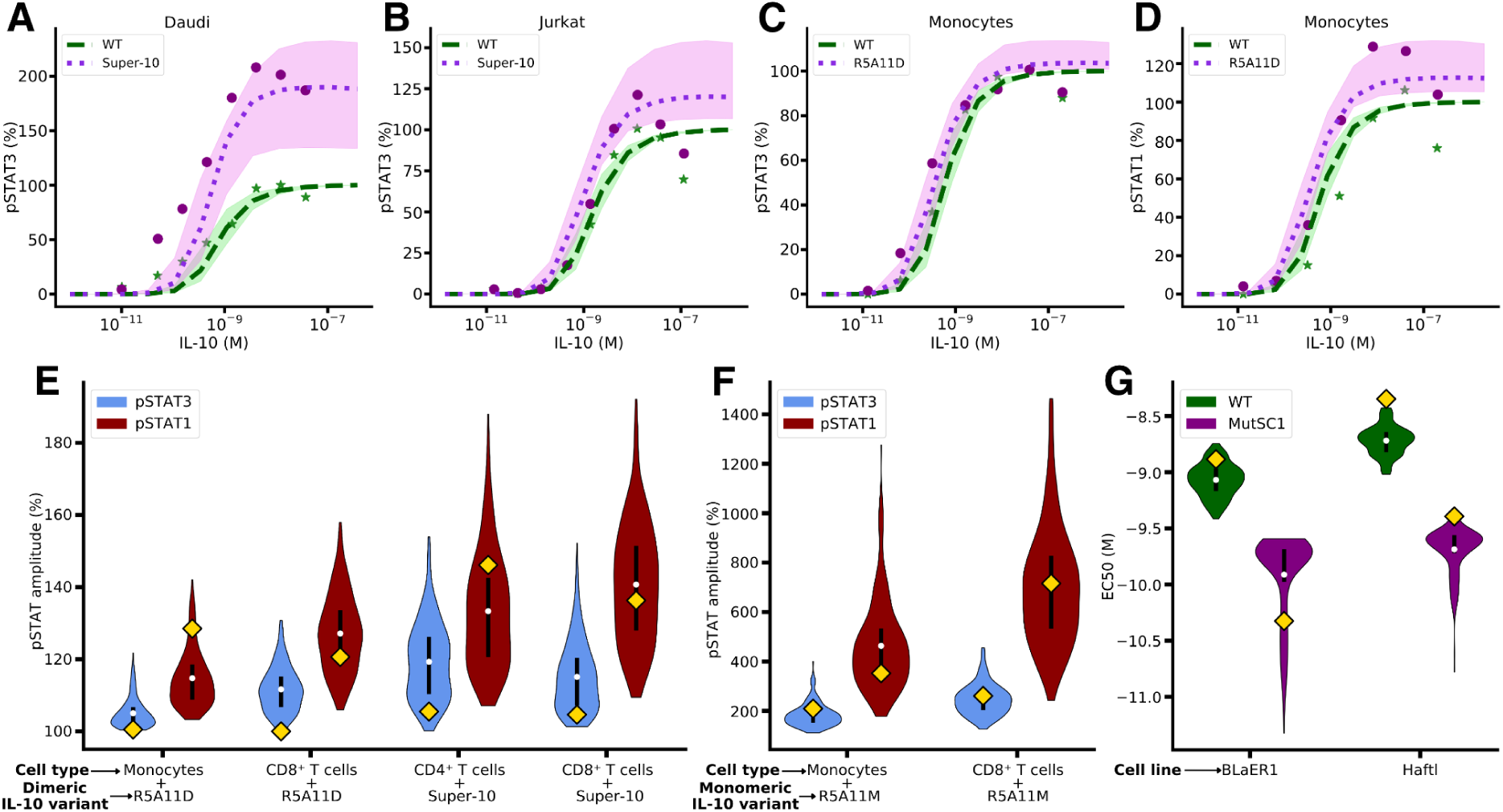
Prediction of in vitro dose-response curves in different cell types and IL-10 mutants. **(A, B)** Dose-response curves of WT IL-10 and Super-10 in the (A) Daudi and (B) Jurkat cell lines for pSTAT3. Model simulations are shown via the discontinuous lines, with the faded area defined by the uncertainty in the initial conditions coming from the RNA to protein linear models. **(C, D)** Dose-response curves of WT IL-10 and R5A11D in monocytes for (C) STAT1 and (D) STAT3. **(E)** Violin plot of the distributions of amplitudes, defined as the maximum minus minimum pSTAT1/pSTAT3 response, of the dose-response curves in the dimeric R5A11D and Super-10 IL-10 mutants across immune cells, assuming uncertainty in the cell-type-specific initial conditions. Yellow squares indicate the experimental values for the dose-response curve’s amplitude. **(F)** Violin plot of the distributions of amplitudes of the dose-response curves in the monomeric R5A11M IL-10 mutant. **(G)** Distribution of EC50s of the dose-response curves in WT IL-10 and the single-chain variant MutSC1 across two cell lines. pSTAT values in all plots are normalized to the maximum WT IL-10 response per cell and displayed as a percentage. IL-10 concentrations and EC50s are shown in moles per liter (*M*).

### Bayesian model selection supports a receptor memory mechanism for STAT1/STAT3 IL-10 signaling

Given the undefined underlying mechanism generating the differential responses of STAT1/STAT3 signaling in IL-10, we formulated three mechanistic models of the IL-10 pathway (Figures 1B, EV1). Given that STAT1 is known not to interact through the STAT3-specific tyrosine residues of IL-10RA, we built a Baseline model in which STAT1 and STAT3 are recruited to IL-10RA through different docking sites that become available only while the IL-10 signaling complex is formed (IL-10+IL-10RA+IL-10RB) (upper row in Figure 1B and Figure EV1A)(Weber-Nordt *et al*, 1996). This attributes STAT1’s greater dependence on IL-10RB to differences in STAT1/STAT3 abundance and binding kinetics. We next defined two simple alternative mechanisms that could account for the observed STAT1/STAT3 asymmetry beyond affinity/expression differences by following prior work on IL-10 and other cytokines (Herrero *et al*, 2003; Riley *et al*, 1999; Rahimi *et al*, 2005; Waters *et al*, 2018). First, we developed a Receptor Memory model (middle row in Figure 1B and Figure EV1B), in which, after the formation of the IL-10 signaling complex, free IL-10RA and IL-10+IL-10RA complexes can signal through STAT3 as long as the IL-10RA-associated Jak1 remains phosphorylated. However, to produce the higher dependence of STAT1 signaling towards IL-10RB binding, STAT1 still needs the IL-10 signaling complex to be phosphorylated at all times, as in the Baseline model. Finally, we built a Kinetic Proofreading model (bottom row in Figure 1B and Figure EV1C) in which STAT1 binds to IL-10RA via an adapter protein (A), whereas STAT3 binds IL-10RA directly, as in the Baseline model.

To determine which mechanism best reproduces the differences between STAT1 and STAT3 seen in experimental dose-response curves around the parameter values reported in the literature, we used the Approximate Bayesian Computation - Sequential Monte Carlo (ABC-SMC) method (Liepe *et al*, 2014; Toni *et al*, 2009). ABC-SMC allows us to obtain parameter distributions and model probabilities constrained by Gaussian priors centered on literature binding rates and fitted to pSTAT1/pSTAT3 dose-response data. As model identifiability represents a concern when dealing with complex parameter inferences, we performed a sloppiness analysis (Gutenkunst *et al*, 2007; Monsalve-Bravo *et al*, 2022) to rank the model’s parameters by their effect on its prediction (Figure EV2). This allows us to limit inference to identifiable parameters, enabling us to run the ABC-SMC algorithm with smaller populations and fewer iterations.

Using this constrained inference framework, we applied ABC-SMC model selection. Under a permissive threshold in the initial iterations, the Receptor Memory and Baseline models hold, while the Kinetic Proofreading model collapses (i.e., the probability that the model explains the data given the parameter priors, becomes *p*_(_*m*|θ_0_, *D*_)_ = 0 ) (Figure 2B). As the error threshold becomes more stringent in later iterations, the Baseline model also collapses. Therefore, model selection by ABC-SMC suggests that a Receptor Memory mechanism could be the driver behind the differences between IL-10-driven STAT3 and STAT1 signaling among the tested hypotheses, but does not exclude other alternative mechanisms. At the same time, ABC-SMC parameter inference yielded narrow, approximately Gaussian posterior distributions (*p*(θ|*D*)) centered within one order of magnitude of their literature values for nearly all parameters of the three IL-10 variants (Figures 2C-F, Appendix Figure S3). Together, these results show that STAT3 Receptor Memory is a plausible mechanistic explanation of the pSTAT1/pSTAT3 dose-response data differences under stringent error thresholds and parameter values near the ones reported in the literature.

### Prediction of variant and cell type-specific IL-10 in vitro responses

By numerically integrating the Receptor Memory model across IL-10 doses, cell types, and IL-10 variants, we used the obtained steady state values to generate predictions on IL-10-pSTAT dose-response curves (Saxton *et al*, 2021; Gorby *et al*, 2020). Using ABC-SMC-inferred parameter values, our model reproduced *in vitro* STAT1/STAT3 responses in a training dataset containing three immune cell types and two cancer cell lines with WT IL-10 and two IL-10 variants with increased IL-10RB binding (Super-10 and R5A11D) (Figure 3A-D, Appendix Figure S4). Performance was quantified using the mean absolute errors (*MAE*) of the concentration at which 50% of the maximum response is achieved (*EC*50) and the amplitude (*AMP*) of dose-response curves (Appendix Table S2). In a test dataset with unseen conditions, the model maintained the goodness-of-fit seen in the training set (Appendix Figures S2C-D, S5). Furthermore, combining the training and test datasets, no statistically significant bias in the model’s performance was detected on any of the simulated cell types or IL-10 mutants (Appendix Figures S2E-H).

As seen in Figure 3E, the Receptor Memory model captured the stronger dependence of STAT1 signaling on IL-10RB binding. IL-10 mutants with increased binding affinity to IL-10RB increased STAT1 phosphorylation at high IL-10 concentrations relative to WT IL-10, while preserving WT levels of STAT3 phosphorylation (Figure 3E). At the same time, even after parameter optimization, the Baseline model failed to reproduce differential STAT1/STAT3 responses and, consistent with ABC-SMC model selection, showed a significantly worse performance in EC50 and amplitude (Figure EV3). This reaffirms that affinity/expression-based differences alone are insufficient to produce differential STAT1/STAT3 responses. At the same time, compared with regression-based model alternatives, the mechanistic Receptor Memory model was the only approach that faithfully reproduced dose-response curves in both training and test datasets (Appendix Table S2).

To further validate our Receptor Memory model, we simulated the response of monomeric (Gorby *et al*, 2020) and single-chain IL-10 variants (Montero-Blay *et al*, 2023). Although WT IL-10 is inactive as a monomer, engineered monomeric IL-10 variants can engage a single IL-10RA and IL-10RB to form a signaling-competent complex with reduced binding affinities (Josephson *et al*, 2000). By restricting the Receptor Memory model to bind a single receptor pair and fine-tuning the IL-10RA dephosphorylation and IL-10RB binding dynamics, we recapitulated the increased differences between pSTAT1 and pSTAT3 signaling in a monomeric IL-10 variant with increased binding affinity to IL-10RB (Figure 3F, Appendix Figure S6). In contrast, single-chain IL-10 dimers retain WT-like binding properties but cannot dissociate into monomers, as they are formed by a single chain of amino acids (Montero-Blay *et al*, 2023). Using the previously inferred WT IL-10 kinetics and blocking IL-10 dissociation, the model reproduced the lower EC50s of single-chain variants in 2 unseen cancer cell lines (Figure 3G).

Overall, the Receptor Memory model showed consistent performance over multiple cell types and IL-10 variants, remaining robust to cell-type differences in receptor expression (Appendix Figure S7). These results support a STAT3-biased Receptor Memory as a mechanistic explanation for the differences in STAT1/STAT3-driven IL-10 signaling, and establish our model as a quantitative method for evaluating how engineered IL-10 variants can reshape IL-10 signaling across cell types.

### From pSTAT dynamics to IL-10-induced gene expression

Although the design of IL-10 variants for therapeutic applications largely centers on modulating STAT1 and STAT3 activation, these phosphorylation patterns provide a simplified picture of IL-10 downstream effects. This is due to the fact that STAT1 and STAT3 phosphorylation are just the initial steps into a gene regulatory program that shapes IL-10 responses across immune cell types (Ouyang & O’Garra, 2019; Saraiva *et al*, 2020; Herrero *et al*, 2003). Therefore, a comprehensive understanding of IL-10 pleiotropy requires integrating IL-10 signaling responses from IL-10 receptor engagement to STAT phosphorylation, together with the downstream changes in gene expression after STAT activation. Having established that the Receptor Memory model accurately recapitulates IL-10-induced STAT1/STAT3 signaling patterns, giving mechanistic insights into how pro-and anti-inflammatory signals emerge in IL-10 at the level of STAT phosphorylation, we next asked how these upstream signaling patterns translate into downstream transcriptional effects.

To assess whether our model encodes information relevant to downstream IL-10 responses, we used single-cell RNA sequencing data (scRNAseq) from mouse lymph nodes (Cui *et al*, 2024, Data ref: Cui *et al*, 2024) to examine whether model-predicted pSTAT1/pSTAT3 abundances correlated with IL-10-induced transcriptomic changes. Due to the high correlation between pSTAT1 and pSTAT3 in the simulated cell types and the fact that signals from both STAT1 and STAT3 responses are combined in the data (Appendix Figure S8), we summed the model-predicted pSTAT1 and pSTAT3 numbers per cell as a readout of IL-10 pathway activation (*pSTAT abundance*). Furthermore, since log2 fold changes are sensitive at low transcript counts and exhibit compressive saturation at high counts (which could distort the underlying shape of the pSTAT-RNA response), we quantified the IL-10 transcriptional response using changes in normalized counts (Δ*RNA counts*). First, we correlated the model-simulated pSTAT1 and pSTAT3 abundances with the mean expression change of IL-10 differentially expressed genes (DEGs) from the immune dictionary 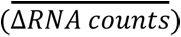 across 11 immune cell types (Figures 4A, Appendix Figure S9)(Cui *et al*, 2024). Although these DEGs were defined in a cell-type-specific manner, we found an approximately linear relationship between pSTAT and the RNA response, which was negative for downregulated genes and positive for upregulated genes (Figure 4A). These results confirm that model predictions of STAT phosphorylation can reflect the scale of the downstream transcriptomic responses.

**Figure 4:**
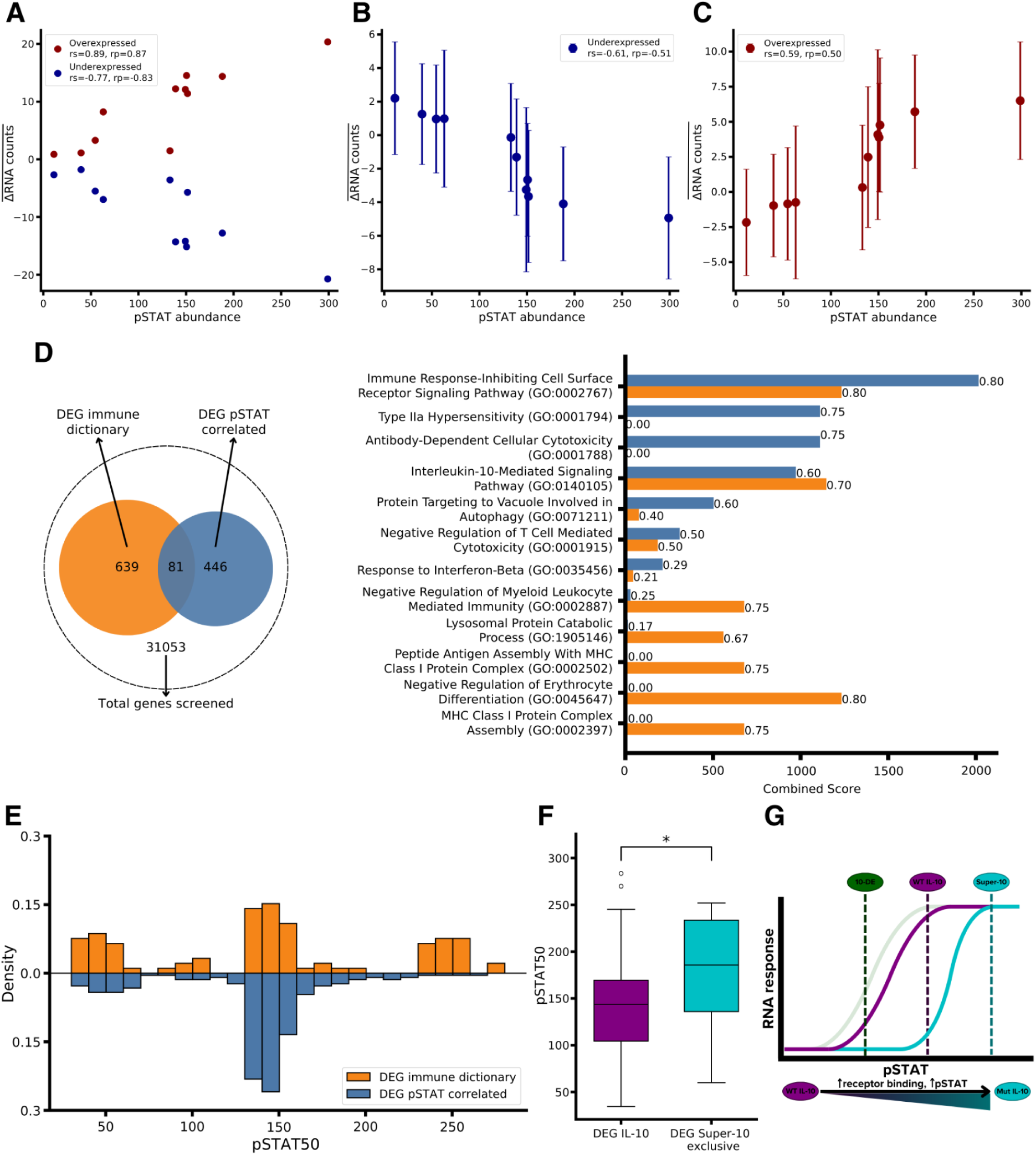
Simulation of in vivo IL-10 mouse response to detect IL-10-regulated genes. **(A)** Scatter plot of the model-simulated pSTAT1 and pSTAT3 in different immune cell types and the mean change in normalized RNA counts in overexpressed and underexpressed gene sets from the Immune dictionary. pSTAT abundance is shown as the number of pSTAT1 and pSTAT3 molecules per cell. **(B, C)** Relationship between model-simulated pSTAT1 and pSTAT3 and the change in RNA counts of genes with (B) strongly negative pSTAT-RNA correlations (underexpressed) or (C) strongly positive (overexpressed) correlations. The points represent the mean, and the error bars represent the standard deviation of the change in RNA counts among genes. **(D)** Overrepresentation analysis (ORA) of DEG lists from the Immune dictionary and the strong pSTAT-RNA correlations. We only show the top 7 significantly enriched sets (Adjusted p-value < 0.05) sorted by combined score for each of the DEG lists. At the end of each bar, we show the percentage of overlap with each gene set. **(E)** Histogram of the distribution of pSTAT activation/repression thresholds (pSTAT50) calculated by fitting logistic curves to gene-specific pSTAT-RNA relationships to the genes in the IL-10 DEG lists from the Immune Dictionary and the strong pSTAT-RNA correlations. **(F)** Box plot of pSTAT50s of genes differentially expressed only by the Super-10 mutant (*N* = 17) or genes differentially expressed also in WT IL-10 (*N* = 49) (*p* = 0. 045). The p-value was calculated using the Scipy implementation of the Mann-Whitney U test. Boxes represent the interquartile range (IQR; Q1–Q3) with the median indicated by a horizontal line while the whiskers extend to the most extreme points within 1.5×IQR. **(G)** Scheme on how IL-10 variants with different signaling strengths could aggregate different transcriptional programs. Variants with reduced receptor engagement (10-DE) produce a reduced version of the WT IL-10-induced transcriptional response. However, a distinct subset of genes becomes activated or repressed only upon stimulation with IL-10 variants with increased STAT phosphorylation (Super-10).

We next aimed to identify additional IL-10-responsive genes based on strong correlations between model-estimated pSTAT abundances and changes in RNA expression. Dose-resolved IL-10 perturbation datasets in a single cell type are not available, preventing a direct evaluation of transcriptomic responses across different pSTAT levels. Nonetheless, pSTAT abundance also varies across cell types at the same IL-10 dose due to differences in receptor expression, effectively providing a range of different IL-10 signaling intensities (Appendix Figure S10). Given the available data, we believe that genes exhibiting strong pSTAT-RNA correlations across cell types are more likely to capture the general IL-10 underlying pSTAT-transcriptome relationship while reducing the influence of other levels of gene regulation, like cell-type-specific effects.

By correlating RNA expression changes per gene (Δ*RNA counts*) to the model-simulated pSTAT abundance in 11 immune cell types, we identified a set of genes induced/repressed by IL-10 in a pSTAT-dependent manner. Separating pSTAT-correlated genes into downregulated and upregulated, we show that these genes produced a mean logistic-like response, with small transcriptomic changes at both low and high pSTAT abundances (Figures 4B-C). Using the IL-10 DEGs from the immune dictionary (639 genes, in orange) as a control and comparing them with the DEGs identified via pSTAT-RNA correlations (446 genes, in blue), we found a low overlap between the 2 groups (81 genes) (Figure 4D)(Cui *et al*, 2024). The poor overlap between DEG lists was also observed in gene-set overrepresentation analysis (ORA), in which the most significantly enriched gene sets were found in only one of the two DEG lists (Figure 4D, Appendix Figure S11). Both lists mostly recovered IL-10-relevant terms, including suppression of immune signaling and regulation of antigen presentation. As a control, pSTAT-RNA-correlated DEGs were recomputed after permuting the Δ*RNA counts* within each gene, causing a total loss of the immune-relevant terms (Appendix Figure S12). At the same time, most of the IL-10-related gene sets emerged from the DEGs common to both lists, rather than the DEGs specific to one of the methodologies (Appendix Figure S13). Taken together, the strong enrichment of immune-related pathways among the pSTAT-RNA-correlated DEGs indicates that these associations reflect an IL-10-linked transcriptional program common across different cell types rather than spurious correlations.

The low overlap between DEG lists could be due to methodological differences. Whereas immune dictionary DEGs are defined independently within each cell type, favoring genes with large expression shifts, our pSTAT-ΔRNA correlation approach integrates information across cell types and can detect IL-10 responsive genes with smaller effects (CD148, IL1RL2) (Appendix Figures S14, S15A-B)(Cui *et al*, 2024). However, genes restricted to a small number (1-3) of cell types (CCL2, CCL7) or regulated at extreme pSTAT levels (SOCS3, DUSP1) are not captured by our correlation-based analysis (Appendix Figure S15C-F). Therefore, both approaches identify complementary modes of IL-10 regulation, including cell-type-specific and shared regulatory programs.

### Regulation at different signaling strengths

IL-10 induces dose- and variant-dependent effects within the same cell type, potentially reflecting the activation of distinct transcriptional programs at different signaling strengths (Naing *et al*, 2018; Jog *et al*, 2018). To test this hypothesis, and based on the logistic-like relationship between pSTAT and transcriptomic changes identified in the previous analysis, and the broader response spectrum captured by the Immune Dictionary DEGs, we fitted logistic models to individual gene-specific pSTAT-RNA relationships. This allowed us to estimate pSTAT abundance thresholds at which each gene is regulated, defined as the model-simulated pSTAT abundance at which each gene reaches 50% of its maximal activation or repression (pSTAT50).

After filtering genes by goodness-of-fit, we compared the distributions of fitted pSTAT50 values in each DEG set (Figure 4E). The distribution of pSTAT thresholds for the correlation-derived DEG set (blue bars in the figure) showed a small peak at low pSTAT abundances and a single dominant peak at intermediate pSTAT levels, consistent with the pattern observed in Figures 4B and 4C, and reflecting the constraints of our methodology. In contrast, the Immune Dictionary DEGs (orange bars in the figure) displayed a distribution with an additional third peak at high pSTAT levels. By clustering the genes from both DEG lists via correlations, we recovered the same low, intermediate, and high pSTAT threshold gene groups (Appendix Figure S16). Finally, overrepresentation analysis showed that these pSTAT-threshold groups were enriched for distinct gene sets (Appendix Figure S17). These results suggest that IL-10 could be aggregating different transcriptional programs engaged at different signaling strengths, partially explaining its pleiotropic behavior.

This idea raises an important consideration for IL-10-based therapies, as IL-10 variants with lower/higher STAT activation could produce distinct biological effects instead of acting as weaker/stronger versions of WT IL-10. To test this hypothesis, we examined whether genes differentially expressed by IL-10 variants with different STAT phosphorylation levels could display shifts in gene expression/repression thresholds. To do that, we analyzed bulk RNA-seq data from human CD8⁺ T cells together with our estimated gene-specific pSTAT50s (Saxton *et al*, 2021; Data ref: Saxton *et al*, 2021). In that work, CD8⁺ T cells were treated at saturating conditions (10 nM) of either WT IL-10, an IL-10 mutant with lower IL-10RB affinity (10-DE), or an IL-10 mutant with higher IL-10RB affinity (Super-10). Calculating fold changes (log2fc) from the resting and IL-10-induced cells, we see that the 10-DE mutant has a global reduction in RNA responses relative to WT IL-10, consistent with its lower STAT phosphorylation (Figure EV4A). However, for genes shared between WT and 10-DE responses, which should therefore require only moderate pSTAT levels for activation/repression, we did not observe a clear shift towards lower pSTAT thresholds (Figure EV4B).

In contrast, a comparison of WT IL-10 with the Super-10 mutant revealed two gene responses that are compatible with its higher STAT phosphorylation: shared genes responsive to WT IL-10 and Super-10 and genes exclusively induced by Super-10 (Figure EV4C). In PCA space, only the Super-10-specific genes separated the Super-10 conditions from PBS, WT, and 10-DE (Figures EV4E-F). Furthermore, Super-10 exclusive genes showed a shift toward higher predicted pSTAT50s, supporting the existence of gene regulation at signaling levels beyond WT IL-10 (Figure 4F). Therefore, most genes seem to be already differentially expressed in response to WT IL-10, whereas a distinct subset becomes activated or repressed only upon stimulation with IL-10 variants that greatly increase STAT phosphorylation (Figure 4G). In contrast, variants with reduced receptor engagement just produced a reduced version of the WT IL-10-induced expression changes rather than qualitatively distinct transcriptional effects.

## Discussion

IL-10 is a key anti-inflammatory cytokine, and compromised IL-10 activity is clearly linked to autoimmune diseases like IBD (Ouyang & O’Garra, 2019; Walter, 2014). At the same time, IL-10 can elicit complex, and often opposing, downstream responses, likely causing the limited disease regression and unexpected adverse effects seen with IL-10-based treatments (Wang *et al*, 2019b; Saxena *et al*, 2015). In this study, we integrated mechanistic models of IL-10 signaling with receptor/STAT expression patterns and Bayesian model selection to help explain IL-10 pleiotropy through differential STAT1/STAT3 activation patterns and downstream transcriptional effects.

Using Bayesian inference, we identified a STAT3-specific mechanism that is consistent with the reduced dependence of STAT3 signaling on IL-10RB binding relative to STAT1 (Gorby *et al*, 2020; Saxton *et al*, 2021). Our Receptor Memory model quantitatively reproduced cell-type and variant-specific pSTAT1/pSTAT3 dose-response curves across multiple immune cell types and cancer cell lines. These results provide a mechanistic explanation for modulating IL-10-based therapies at the level of STAT phosphorylation toward pSTAT1-mediated pro-inflammatory responses or pSTAT3-mediated anti-inflammatory responses through changes in IL-10RB binding. Furthermore, our model generalized for distinct IL-10 variants not used in model inference, including monomeric and single-chain constructs, establishing it as an in silico platform to predict how novel IL-10 variants could reshape IL-10 signaling.

However, to reproduce experimental dose-response curves, our IL-10 signaling model underestimated the experimentally reported differences in IL-10RB binding between WT IL-10 and variants with enhanced IL-10RB affinity (R5A11 and Super-10)(Gorby *et al*, 2020; Saxton *et al*, 2021; Yoon *et al*, 2005; Logsdon *et al*, 2002). These experimentally measured binding affinities are difficult to reconcile simultaneously with the modest differences observed in pSTAT3 signaling between WT IL-10 and the IL-10RB-enhanced variants, and with the measured expression levels of IL-10RA and IL-10RB across immune cell types. This suggests that either one or more of these measurements might deviate beyond a reasonable experimental uncertainty, or that an additional mechanism, such as receptor binding memory, buffers STAT3 activation against variations in IL-10RB affinity.

The discrepancy between predicted and measured binding rates could also reflect differences between receptor interactions in Biacore-based measurements and at the cell surface. For example, receptors are immobilized on a surface in Biacore assays, whereas they can freely diffuse in the plasma membrane (Gorby *et al*, 2020). In addition, Biacore experiments typically use monomeric IL-10 to simplify interactions, despite the reduced affinity of monomeric forms compared to the dimeric WT IL-10 (Gorby *et al*, 2020; Saxton *et al*, 2021). Although these experimental constraints could bias the apparent binding kinetics, our inferred binding kinetics still remained consistent with literature values within an order of magnitude.

A similar issue emerged when modeling the decline in signaling observed at high IL-10 concentrations, as the behavior could be universally reproduced only by assuming a binding of IL-10RB to free IL-10 that substantially exceeds published experimental values (Yoon *et al*, 2005; Logsdon *et al*, 2002). This is because high IL-10RB binding rates allow the sequestration of IL-10RB by free IL-10 at high IL-10 doses, limiting the formation of complexes with both receptors and reducing the IL-10 signaling response. An alternative explanation for the reduced signaling at elevated ligand concentrations could be the existence of negative feedback mechanisms like IL-10RA degradation via ubiquitination, or IL-10-derived changes in the expression of IL-10 receptors or cytokine signaling inhibitors like SOCS3 (Ding *et al*, 2003; Jiang *et al*, 2011; Cui *et al*, 2024). These mechanisms would limit IL-10RA availability under high signaling conditions, thereby affecting both STAT1 and STAT3 signaling in a manner consistent with experimentally measured dose-response curves. Using the same ABC-SMC model selection framework, these hypotheses could also be tested in silico.

Having confirmed that our IL-10 model encodes information relevant to downstream RNA responses, we explored how pleiotropic responses emerge at the transcriptomic level in IL-10. By correlating cell-type-specific pSTAT model predictions with downstream transcriptomic changes, we identified a set of IL-10-responsive genes that reflect a shared, immune regulatory program and follow a mean logistic pSTAT-RNA relationship. At the same time, we demonstrated how IL-10 could aggregate different transcriptional programs engaged at different signaling strengths. This creates a more complex picture of IL-10-derived transcriptional regulation, where a fixed set of downstream effects is not linearly dependent on IL-10 signaling strength. Instead, when increasing IL-10 pathway activation, we might find totally different downstream effects derived from distinct genes being activated/inactivated in a nonlinear way at increasing IL-10 signaling strengths. This could therefore explain the emergence of IL-10 dose-specific effects like CD8⁺ T cell proliferation and NK cell cytotoxicity at high IL-10 concentrations or how the viral IL-10 drives an enhanced immune silencing when compared to the WT IL-10 (Ouyang & O’Garra, 2019; Wang *et al*, 2021; Jog *et al*, 2018).

Furthermore, these results reveal an important limitation of using pSTAT dose-response curves to validate novel IL-10-based therapies. First, saturation implies that IL-10 variants producing higher STAT1/3 phosphorylation might not necessarily translate into further changes in downstream effects for genes with low-threshold IL-10 responses. At the same time, these IL-10 variants could trigger additional unexpected effects over wild-type IL-10 for genes with high pSTAT thresholds. Accordingly, engineering IL-10 variants to modify the maximum response may produce qualitatively distinct and potentially unanticipated transcriptional responses relative to WT IL-10 (Figure EV5). In contrast, mutations that primarily shift the EC50 of the IL-10-pSTAT dose-response relationship are expected to yield WT-like RNA responses, but at different ligand concentrations (Figure EV5). All in all, these findings highlight the need to guide novel IL-10-based therapy design with downstream functional readouts, enabling the early prediction of potentially unexpected effects.

Still, our predictive capacity of these effects could be constrained by the available data, which does not allow us to capture cell-type-specific effects. This limitation could be addressed in future studies with new data collected across multiple IL-10 concentrations rather than a single dose. This would enable not only the identification of cell-type-specific differentially expressed genes correlated with pSTAT, but would also give a more comprehensive understanding of how increasing IL-10 doses, and corresponding levels of STAT phosphorylation, reshape downstream transcriptomic changes across distinct immune cell types. Furthermore, in this study, we combined STAT1 and STAT3 predictions as a measure of IL-10 pathway activation to determine IL-10 downstream transcriptional responses. However, given IL-10 perturbation data in STAT1 or STAT3-deficient cells, we could also begin to decouple STAT1 and STAT3-driven IL-10 responses at the transcriptomic level.

In conclusion, our study highlights the potential of computational modeling as a powerful framework to refine our understanding of the IL-10 pathway and to begin disentangling the pleiotropic effects of IL-10 from receptor binding kinetics to downstream transcriptional responses, with the future goal of supporting the rational development and clinical translation of more precise and effective IL-10-based therapies.

## Methods

### Quantification of IL-10 receptors

HEK-Blue IL-10 cells (hkb-il10v2_Invivogen) were cultured in DMEM-10% FBS. When reaching 90% of confluence, cells were washed with PBS and lysed in 100 μl of Urea 6 M, NH_4_HCO_3_ 0.2 M. The samples were sonicated for 10 min at high power with 30-second on/off pulses (Diagenode Bioruptor), then centrifuged at 4 °C for 10 min at 16000 g. Total protein in the extracts was determined by BCA assay (Pierce) and adjusted to 1 μg protein/μl. Samples (10 µg) were reduced with dithiothreitol (30 nmol, 37 °C, 60 min) and alkylated in the dark with iodoacetamide (60 nmol, 25 °C, 30 min). The resulting protein extract was first diluted to 2M urea with 200 mM ammonium bicarbonate for digestion with endoproteinase LysC (1:10 w:w, 37°C, over 6h, Wako, cat # 129-02541) and then diluted 2-fold with 200 mM ammonium bicarbonate for trypsin digestion (1:10 w:w, 37°C, over 8h, Promega cat # V5113). After digestion, the peptide mix was acidified with formic acid and desalted with a MicroSpin C18 column (The Nest Group, Inc) before LC-MS/MS analysis.

Samples were analyzed using an Orbitrap Fusion Lumos mass spectrometer (Thermo Fisher Scientific, San Jose, CA, USA) coupled to an EASY-nLC 1200 (Thermo Fisher Scientific (Proxeon), Odense, Denmark). Peptides were loaded directly onto the analytical column and were separated by reversed-phase chromatography using a 50-cm column with an inner diameter of 75 μm, packed with 2 μm C18 particles (Thermo Fisher Scientific, cat # ES903). Chromatographic gradients started at 95% buffer A and 5% buffer B with a flow rate of 300 nl/min and gradually increased to 25% buffer B and 75% A in 79 min and then to 40% buffer B and 60% A in 11 min. After each analysis, the column was washed for 10 min with 100% buffer B. Buffer A: 0.1% formic acid in water. Buffer B: 0.1% formic acid in 80% acetonitrile. The mass spectrometer was operated in positive ionization mode with nanospray voltage set at 2.4 kV and source temperature at 305°C.

To identify suitable peptides for absolute quantification of IL-10RA and IL-10RB, label-free proteomic analysis was performed on the protein extract. The acquisition was performed in data-dependent acquisition (DDA) mode, and full MS scans with 1 micro scan at a resolution of 120,000 were used over a mass range of m/z 350-1400 with detection in the Orbitrap mass analyzer. Auto gain control (AGC) was set to ‘standard’ and injection time to ‘auto’. In each cycle of data-dependent acquisition analysis, following each survey scan, the most intense ions above a threshold ion count of 10000 were selected for fragmentation. The number of selected precursor ions for fragmentation was determined by the “Top Speed” acquisition algorithm and a dynamic exclusion of 60 seconds. Fragment ion spectra were produced via high-energy collision dissociation (HCD) at a normalized collision energy of 28%, and they were acquired in the ion trap mass analyzer. AGC was set to 2E4, and an isolation window of 0.7 m/z and a maximum injection time of 12 ms were used. Digested bovine serum albumin (New England Biolabs, P8108S) was analyzed between each sample to avoid sample carryover and to ensure stability of the instrument, and QCloud has been used to control instrument longitudinal performance during the project (Chiva *et al*, 2018).

Acquired spectra were analyzed using the Proteome Discoverer software suite (v2.5, Thermo Fisher Scientific) and the Mascot search engine (v2.6, Matrix Science)(Perkins *et al*, 1999). The data were searched against a Swiss-Prot human database (as of April 2022, 20376 entries) plus a list of common contaminants and all the corresponding decoy entries (Beer *et al*, 2017). For peptide identification, a precursor ion mass tolerance of 7 ppm was used for the MS1 level, trypsin was chosen as the enzyme, and up to three missed cleavages were allowed. The fragment ion mass tolerance was set to 0.5 Da for MS2 spectra. Oxidation of methionine and N-terminal protein acetylation were used as variable modifications, whereas carbamidomethylation on cysteines was set as a fixed modification. False discovery rate (FDR) in peptide identification was set to a maximum of 5%. Peptides corresponding to IL-10RA and IL-10RB were evaluated based on highly observable, signal intensity,y and uniqueness to the target proteins (Desiere *et al*, 2006). The selected peptides (IL-10RA (Q13651): EECISLTR, TGCLEEESPLTDGLGPK, GQDDSGIDLVQNSEGR, and IL-10RB (Q08334): LSVIAEDSESGK, IENEYETWTMK, FQITPQYDFEVLR) were chemically synthesized as AQUA Ultimate grade using C-terminal 13C6, 15N2-Lysine or 13C6, 15N4-Arginine (Thermo Fisher Scientific, Germany).

For absolute quantification of IL-10 receptors, we cultured U-937, HuT 78, THP-1, and K-562 cell lines in RPMI-1640 media-10% FBS. Protein extracts from the cultures were extracted in the same way as the HEK-Blue IL-10 cells. 3 µg of protein extract were analyzed together with 0.075 pmol of heavy isotope–labeled peptides corresponding to IL-10RA and IL-10RB using a parallel reaction monitoring (PRM) method. A full MS scan at a resolution of 30,000 was used over a mass range of m/z 350-1400 with detection in the Orbitrap mass analyzer. For the PRM data acquisition, a quadrupole isolation window is set to 1.4 m/z, and MSMS scans over a mass range of m/z 300-2000, with detection in the Orbitrap at a resolution of 60,000. MSMS fragmentation was performed using HCD at 30 NCE, the auto gain control (AGC) was set to 1e5, and a maximum injection time of 118 ms. The m/z values corresponding to the doubly charged light and heavy forms of the selected peptide were defined in the inclusion mass list for subsequent fragmentation. Skyline software (26.0.9.032) was used to extract the fragment areas of each peptide and the fold change ratio relative to the internal standard (MacLean *et al*, 2010). Finally, absolute quantification was performed using isotope-labeled peptides as internal standards, as described in Picotti et. al.’s work, and converted to copies per cell based on Avogadro’s number and cell counts (Picotti & Aebersold, 2012). The total number of cells in the extract was estimated via Countess Cell Counting Chamber Slides and an Invitrogen Countess 3 FL Automated Cell Counter. All the raw proteomics data have been deposited in the PRIDE repository with the dataset identifier PXD076154 (Perez-Riverol *et al*, 2022).

### Gene-specific linear models

Inferring protein abundance from mRNA measurements allows us to bypass the limited availability of absolute protein measurements. Although global transcriptome-proteome correlations are typically weak, predictions can be improved by modeling protein copy number as a gene-specific function of transcript levels (Crick, 1958; Edfors *et al*, 2016; Wang *et al*, 2019a; Weber *et al*, 2023). This reflects gene-dependent translation efficiencies and protein half-lives, which are typically conserved across cell and tissue contexts (Edfors *et al*, 2016). On this basis, we developed gene-specific linear models for 4 proteins present in the IL-10 pathway: IL-10RA, IL-10RB, STAT1, and STAT3. These models were trained using the TheilSenRegressor function in the scikit-learn package on bulk RNA-seq data normalized by TPM (transcripts per million) and absolute proteomic quantifications from peripheral blood mononuclear cells and cancer cell lines (Appendix Table S6)(Sikkema *et al*; Rieckmann *et al*, 2017; von Haehling *et al*, 2015; Aizaki *et al*, 2021; Jurlander *et al*, 1997; Tian *et al*, 2014; Data ref: Barretina *et al*, 2019; Buitinck *et al*, 2013; Barretina *et al*, 2019; Sikkema *et al*, 2023). Linear models were tested with transcriptomic and proteomic data from 29 healthy human tissues (Eraslan *et al*, 2019).

### Model Building with PySB

IL-10 signaling models were implemented in PySB (Python 3.10), a rule-based framework for constructing ordinary differential equation (ODE) models of signal-transduction networks (Lopez *et al*, 2013). As all models represent IL-10 signaling in a single cell, 3 PySB compartments were defined: extracellular space, plasma membrane, and cytoplasm. All models’ equations were derived by mass-action laws, where the rate of each reaction is proportional to the product of the concentrations of the interacting species (An Introduction to Systems Biology: Design Principles of Biological Circuits. Second Edition. By Uri Alon. A Chapman & Hall Book. Boca Raton (Florida): CRC Press (Taylor & Francis Group). 143.96 (hardcover); 55.96 (paper). xviii + 324 p.; ill.; index. ISBN: 978-1-138-49011-6 (hc); 978-1-4398-3717-7 (pb). 2020, 2021). As reactions are simulated in molecule-number units, all binding rates were converted from 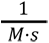 to 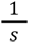 by multiplying them by the Avogadro number (*N*_a_) and the volume of the compartment (*V*_c_) where the resulting protein complex is not anchored to (Equation 1)(Sekar & Faeder, 2018). For reactions happening at the plasma membrane, the binding rate is instead divided by an effective volume that comes from multiplying the cell’s surface area by the plasma membrane’s width (Sekar & Faeder, 2018; Chylek *et al*, 2015).

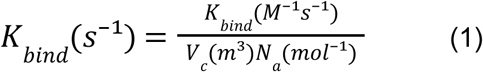

Because the mechanism behind differential STAT1/STAT3 signaling of IL-10 remains unclear, we developed three mechanistic models of the IL-10 pathway (Baseline, Receptor Memory, and Kinetic proofreading). To help visualize the reactions in all 3 models, we generated a schematic depiction of the interacting molecular species and the biochemical reactions that define the 3 models (Figure EV1).

All three models considered IL-10 monomer dimerization in the extracellular space. We also did not assume a specific order of receptor engagement to the IL-10 dimers. Therefore, all receptor binding reactions that formed protein complexes that are a combination of a single IL-10 dimer and a maximum of two IL-10RA and/or two IL-10RB copies were considered in all models, even though direct binding of IL-10 to IL-10RB is known to be weak (Logsdon *et al*, 2002; Yoon *et al*, 2005). Since IL-10RA is known to improve IL-10RB binding, a separate value for its binding/unbinding rate to the IL-10+IL-10RA complex is also considered (K*_IL-RA-RB-b_*) (Logsdon *et al*, 2002; Gorby *et al*, 2020). This is not the case for the binding of IL-10RA to an IL-10+IL-10RB complex, where the models use the binding rate of free IL-10 to IL-10RA (K*_IL-RA-b_*), but scaled to the cell surface’s effective volume.

In the intracellular compartment, all 3 models represent different ideas on how STAT3 signaling can be buffered against variations in IL-10RB affinity with respect to the more variable STAT1 (Saxton *et al*, 2021; Gorby *et al*, 2020). In the Baseline model, any complex in which both IL-10RA and IL-10RB occupy at least one subunit of the IL-10 dimer is signaling-competent and can recruit unphosphorylated STAT1 and STAT3 through its IL-10RA, with one separate binding site per STAT (Figure 1B and EV1A) (Weber-Nordt *et al*, 1996). Upon binding, STAT1/3 can either dissociate at a fixed rate or when the signaling complex is broken. Furthermore, bound STAT1/3 is phosphorylated at a constant rate and released immediately after. Under these assumptions, STAT1’s higher apparent IL-10RB dependence could arise from differences in STAT1/3 abundance and binding/unbinding kinetics. For the Receptor Memory model, STAT1 interacted with IL-10RA in the same way as the Baseline model. Instead, STAT3 could interact with IL-10RA as long as the IL-10RA-associated Jak1 remained phosphorylated (Figure 1B and EV1B). As the receptor-associated kinases are only phosphorylated when the full IL-10+IL-10RA+IL-10RB signaling complex is formed, STAT3 signaling was still dependent on IL-10RB binding, but to a lesser extent than STAT1. IL-10RA copies that are not bound to IL-10 and IL-10RB at the same time are dephosphorylated at a constant rate, thus preventing STAT3 binding and phosphorylation. Finally, in the Kinetic Proofreading model, STAT3 interacted with IL-10RA as in the Baseline model. However, STAT1 interaction with IL-10RA was dependent on the previous binding of an adapter protein, which binds only to the full IL-10+IL-10RA+IL-10RB signaling complex (Figure 1B and EV1C). This generated a higher dependence for IL-10RB on STAT1 signaling, when compared to STAT3. For all models, free pSTAT1/3 was dephosphorylated at a constant rate. The simplified interaction networks, parameter values, and explanations of the models’ parameters can be found in Figure EV1 and Appendix Tables S3 and S4. In the appendix, we also included the resulting ODE system of the Receptor Memory model (Appendix Equations S1-S29).

To speed up simulations during model selection, we derived the steady-state solution of the three models using the linear framework (Nam *et al*, 2022; Ahsendorf *et al*, 2014). As the linear framework has a limit on the model’s complexity, all 3 models needed to be simplified. For the Baseline and Receptor Memory model, STAT1 and STAT3 binding, phosphorylation, and dephosphorylation were considered to be in a quasi-steady state due to their low effect on the model’s response (Figures EV2A, B) (see Appendix for the derivation of the pSTAT1/3 quasi-steady equations). For the Receptor memory and Kinetic Proofreading models, IL-10RB was assumed to only bind after IL-10RA binding, as the binding of IL-10RB to free IL-10 is very low (Logsdon *et al*, 2002; Gorby *et al*, 2020). Since these assumptions introduced deviations from the full ODE formulations that increased the error in the EC50 of the Receptor Memory model, all subsequent simulations and analyses following model selection were performed instead using the original ODE-based models (Appendix Table S3).

### Sloppiness analysis

Parameter inference often yields high uncertainty for parameters that have little influence on model outputs (sloppy parameters), whereas parameters that strongly shape the response (stiff parameters) must be constrained more tightly to not jeopardize the model’s fit (Monsalve-Bravo *et al*, 2022; Gutenkunst *et al*, 2007). To help identify stiff parameters and optimize the subsequent parameter inference, we performed a sloppiness analysis on all parameters of the three models.

By computing the Levenberg-Marquardt Hessian matrix of the model’s cost function (*H*_ij_), we assessed the contribution of parameter combinations to the model’s fit via eigendecomposition (Equation 2)(Monsalve-Bravo *et al*, 2022). To evaluate the model’s fit, we used as a cost function the weighted sum of squares of the error on the IL-10 concentration producing 50% of the maximal response (*EC*50) and amplitude (*AMP*) of the dose-response curve of every variant/cell type pair in our training set (Equation 3). To balance the error coming from the EC50 and the amplitude, the cost function was normalized with two weights (*w_EC_*_50_,*w_AMP_*) that were calculated with the initial literature parameter value set 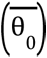 (Equations 4-5). This cost function avoids bias from non-uniform sampling of IL-10 concentrations, where pointwise fitting would overweight densely sampled regions and overfit baseline or saturation regimes while still capturing the global shape of the dose-response relationship.

The evaluated dataset included data on WT IL-10 and 2 mutants with increased IL-10RB binding (R5A11D and Super-10) in 3 immune cell types (Monocytes, CD8⁺, and CD4⁺ T cells) and 2 cancer cell lines (Daudi and THP-1)(Saxton *et al*, 2021; Gorby *et al*, 2020). For each dose-response curve, cells were stimulated with 25 evenly spaced IL-10 doses spanning the minimum and maximum concentrations used in the corresponding in vitro data. Amplitudes and EC50s were obtained by fitting a Hill function to both experimental dose-response data (*EC*50*_obs_*,*AMP_obs_*) and simulated curves 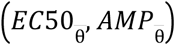 using SciPy’s curve_fit function (Figure EV2)(Virtanen *et al*, 2020).

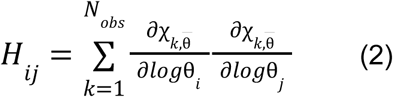

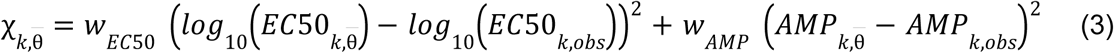

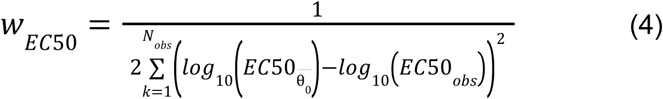

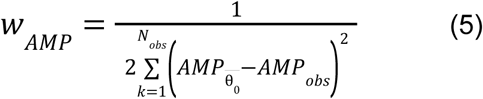

### Bayesian inference and model selection

Bayesian inference allows us to obtain posterior parameter distributions (*p*(θ|*D*)) and model (*m*) probabilities (*p*(*m*|θ_0_, *D*)), conditioned on prior knowledge of the model’s parameters (θ_0_) and experimental data to fit the models (*D*). Approximate Bayesian Computation-Sequential Monte Carlo (ABC-SMC) makes Bayesian inference possible for problems with intractable likelihoods by iteratively approximating posterior probabilities (Liepe *et al*, 2014; Toni *et al*, 2009).

Model selection and parameter estimation were done simultaneously using the pyABC implementation of ABC-SMC, with the MulticoreEvalParallel sampler and an adaptive population strategy that started at 750 individuals and had a limit of 1250. Prior distributions of the parameters were assumed to be normally distributed, centred on literature mean values, and with a constant variance that excluded values beyond 1 magnitude of the experimental data (σ^2^ = 0. 4375). Initial parameter values describing IL-10 and receptor binding events were obtained from in vitro experiments, while STAT1/3 binding, phosphorylation, and dephosphorylation parameters were taken from previously established models (Appendix Tables S3-4)(Faeder *et al*, 2009; Ballas, 1987; Powell *et al*, 2003; Das & Jayaprakash, 2021; Krombach *et al*, 1997; Zhao *et al*, 2008; Syto *et al*, 1998; Logsdon *et al*, 2002; Yoon *et al*, 2005, 2012; Weber-Nordt *et al*, 1996; Wilmes *et al*, 2021; Schindler *et al*, 1995; Reeh *et al*, 2019; Saxton *et al*, 2021; Gorby *et al*, 2020; Maiti *et al*, 2014). Soft bounds were imposed by adding a penalty to the cost function when parameter values deviated by more than one order of magnitude from their literature mean.

Given the results on parameter identifiability coming from the previously performed sloppiness analysis, we fitted the IL-10 variant-inespecific parameters of the cell’s surface (*S_cell_*), IL-10 dimerization unbinding rate (*K_ILM-b_*), and unbinding rate of IL-10RA to IL-10 (*K_IL-RA-b_*) for all three models. For the unbinding rate of IL-10RB to IL-10+IL-10RA (*K_IL-RA-RB-b_*), we fitted each IL-10 variant to have its own parameter value. We additionally fitted the dephosphorylation rate of the receptor (*K_DEPHOS-R_*) for the Receptor Memory model. For the Kinetic Proofreading model, we also estimated the binding rate of STAT1 to the adapter protein (*K_STAT1-f_*), the binding rate of the adapter protein to IL-10RA (*K_A-f_*), and the initial quantity of adapter protein per cell (*A*_0_).

Model selection was performed using the same cost function and training dataset as the sloppiness analysis, but with 6 measurements per dose-response curve, which were centered on the experimentally observed EC50s instead. Given the assumptions made when computing the steady-state approximations of the Receptor Memory model, the model was later recalibrated using ODE simulations with the previously stated ABC-SMC hyperparameters to the same training set used in model selection to ensure an optimal fit. Given differences between WT IL-10 and its monomeric signaling-capable variants, IL-10RA dephosphorylation and IL-10RB binding were inferred again using data on signaling-capable monomeric IL-10 variants. Both parameter estimations used the same described hyperparameter settings as the model selection.

### Simulation of in vitro dose-response curves

After integrating all bulk RNA-seq and absolute proteomics measurements into a single dataset, simulations of in vitro dose-response curves across immune cell types and cancer cell lines were performed with either real protein count measurements or estimations coming from linear models if experimental data were not available (Appendix Table S6)(Sikkema *et al*; Rieckmann *et al*, 2017; von Haehling *et al*, 2015; Aizaki *et al*, 2021; Jurlander *et al*, 1997; Tian *et al*, 2014; Data ref: Barretina *et al*, 2019; Zhang *et al*, 2017; Gaidt *et al*, 2021; Wang *et al*, 2023; Barretina *et al*, 2019; Sikkema *et al*, 2023; Data ref: Gaidt *et al*, 2021; Data ref: Wang *et al*, 2023). Parameter values for all IL-10 variants were extracted from the peak of the ABC-SMC-derived posterior distributions. For the signaling-capable monomeric variants of IL-10, we blocked IL-10 dimerization and only allowed the binding of the receptors on one side of an IL-10 molecule (Josephson *et al*, 2000). For the single-chain IL-10 variants, we instead used the WT IL-10 binding rates, but blocked IL-10 dimer dissociation (Montero-Blay *et al*, 2023). Each dose-response curve was simulated using 24 evenly spaced IL-10 doses. Uncertainty in the initial conditions from the RNA to protein linear models was considered in the dose-response curves by sampling from a normal distribution centred on the linear model’s protein count estimation and then simulating the IL-10 response. The standard deviation of the normal distribution was taken from the mean absolute error of the RNA to protein linear models (σ = 0. 2).

### Prediction of IL-10 gene expression from pSTAT estimations

To predict IL-10 transcriptional programs, we generated pseudo-bulk data from Cui et al.’s single-cell RNA sequencing cytokine perturbation data on immune cells of mouse lymph nodes (Cui *et al*, 2024, Data ref: Cui *et al*, 2024). Pseudobulking was performed by aggregating counts and normalizing them as counts per million (CPM) for each non-stimulated cell type, allowing us to estimate the receptor/STAT protein counts. Given the fixed IL-10 concentration used in the same study (2. 43·10^−6^), we estimated the STAT1/STAT3 phosphorylation after IL-10 induction for each of the assayed cell types using our IL-10 signaling model. To measure the downstream effect of IL-10 on the transcriptome, we pseudobulked all control and IL-10-stimulated cell types, removing cell types with fewer than 40 single cells and genes with less than 10 total reads. After that, we normalized the counts using PyDeseq2’s median of ratios method to perform library-size normalization while minimizing the loss of signal (Muzellec *et al*, 2023). Finally, to measure changes in the transcriptome, we calculated the difference between the normalized counts of resting and IL-10 perturbed cells per gene per cell type (Δ*RNA counts*).

To evaluate whether the model captures information relevant to downstream IL-10 responses, a curated list of IL-10 DEGs per cell type was extracted from the Immune Dictionary, separating the genes into overexpressed (*log*_2_*fc* > 0) and underexpressed (*log*_2_*fc* < 0) (Cui et al, 2024). Given the high correlation between the simulated STAT1 and STAT3 in these cell types, and that both responses are combined in the RNAseq data, we added the STAT1 and STAT3 model estimations to serve as a measure of IL-10 pathway activation (*pSTAT abundance*). Spearman and Pearson correlations between the mean expression of overexpressed and underexpressed DEG from the Immune dictionary were calculated using their SciPy implementation (Virtanen *et al*, 2020). To determine IL-10 DEGs that were dependent on pSTAT, we correlated the expression change per gene (Δ*RNA counts*) across 11 immune cell types to the model-simulated pSTAT via Spearman’s correlation (*r*_s_), filtering by significance (*p* < 0. 05), and dividing them into overexpressed 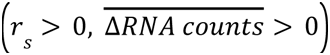 and underexpressed genes 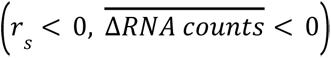. By filtering both by direction (*r*_s_) and mean Δ*RNA counts* across cell types, we only keep genes that have a positive or negative relationship with pSTAT and whose expression is increased or decreased, respectively. As a control, this analysis was repeated, but permuting the Δ*RNA counts* within each gene. Gene set overrepresentation analysis (ORA) was performed on both DEG lists using GSEApy’s Enrichr interface with the GO_Biological_Process_2025 and GO_Molecular_Function_2025 libraries (Fang *et al*, 2023).

### Prediction of IL-10 regulation at different pSTAT thresholds

Based on the logistic-like relationship between pSTAT levels and the mean RNA expression identified in our correlation analysis, we fitted a logistic function to each pSTAT-ΔRNA scatter plot per gene using SciPy’s curve_fit, allowing us to compute gene-specific activation/repression thresholds (pSTAT50) for the downstream RNA response (Equation 6)(Virtanen *et al*, 2020). The function also considers a basal level of change in RNA expression (Δ*counts*_0_), a sharpness of the response (*k*) and an amplitude of the curve (*L*).

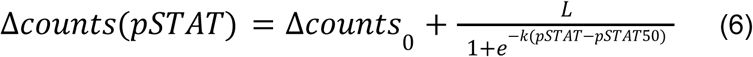

To independently demonstrate IL-10 regulation at different pSTAT thresholds, we clustered genes based on the similarity of the shape of their RNA response. Gene-to-gene distances were calculated as 1 − *abs*(*PearsonR*), and hierarchical clustering was performed with SciPy’s function linkage. Clusters were defined by cutting the dendrogram into four groups using the function fcluster. Furthermore, gene set overrepresentation analysis (ORA) was performed on the DEGs of both lists (Immune dictionary and strong pSTAT-ΔRNA correlations), filtered into genes with low pSTAT thresholds (120<pSTAT50), genes with medium pSTAT thresholds (120<pSTAT50<180), and genes with high pSTAT thresholds (pSTAT50>180). These groupings were decided to compensate for the fact that the majority of genes are found in the medium pSTAT threshold range.

Finally, we used bulk RNA-seq data from CD8⁺ T cells to compare the response between IL-10 variants with different signaling strengths (Saxton *et al*, 2021; Data ref: Saxton *et al*, 2021). Log2 fold changes (*log*2*fc*) between PBS and IL-10 variants were calculated using PyDeseq2, while principal component analysis was performed with the package scikit-learn (Muzellec *et al*, 2023; Buitinck *et al*, 2013). To remove noise, we restricted the analysis to genes activated by at least one of the IL-10 variants (DEG IL-10), which were defined by filtering by effect size (|*log*2*fc*| > 0. 75) and significance (*p_adjusted_* < 0.05) . Genes exclusively differentially expressed in Super-10 (DEG Super-10) were defined as genes with low expression changes in WT IL-10 (|*log*2*fc*| < 1. 25) and having 33% times higher log2fc in Super-10 than in WT IL-10. Given the very low effect of the 10-DE variant, we defined genes differentially expressed in 10-DE (DEG 10-DE) as genes whose log2 fold change in 10-DE is no more than 2.5-fold lower than in WT IL-10. This allowed us to select genes with similar changes in expression between PBS and WT IL-10 and PBS and 10-DE conditions.

## Expanded view figure legends

**Figure EV1:**
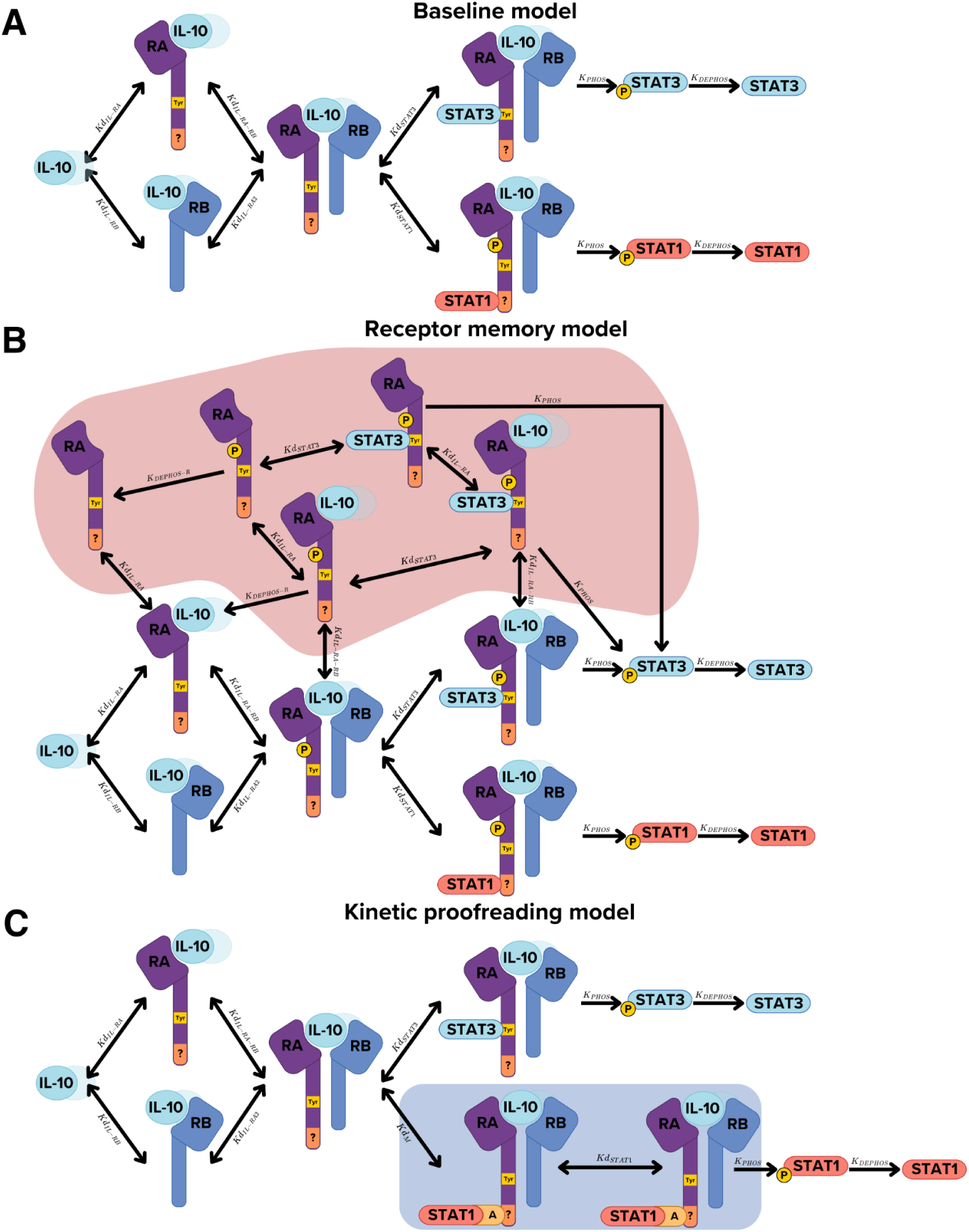
Interaction networks for IL-10 pathway models. **(A-C)** Interaction network of the (A) Baseline, (B) Receptor Memory, and (C) Kinetic Proofreading models. For visualization purposes, the interaction networks shown here were simplified relative to the full model implementations. First, we show receptor binding to only one of the IL-10 subunits in the dimer. Furthermore, in all models, STAT1 and STAT3 binding can happen simultaneously. This includes the adapter protein A for the Kinetic Proofreading model. The red and blue shades represent interactions that differ from those in the Baseline model.

**Figure EV2:**
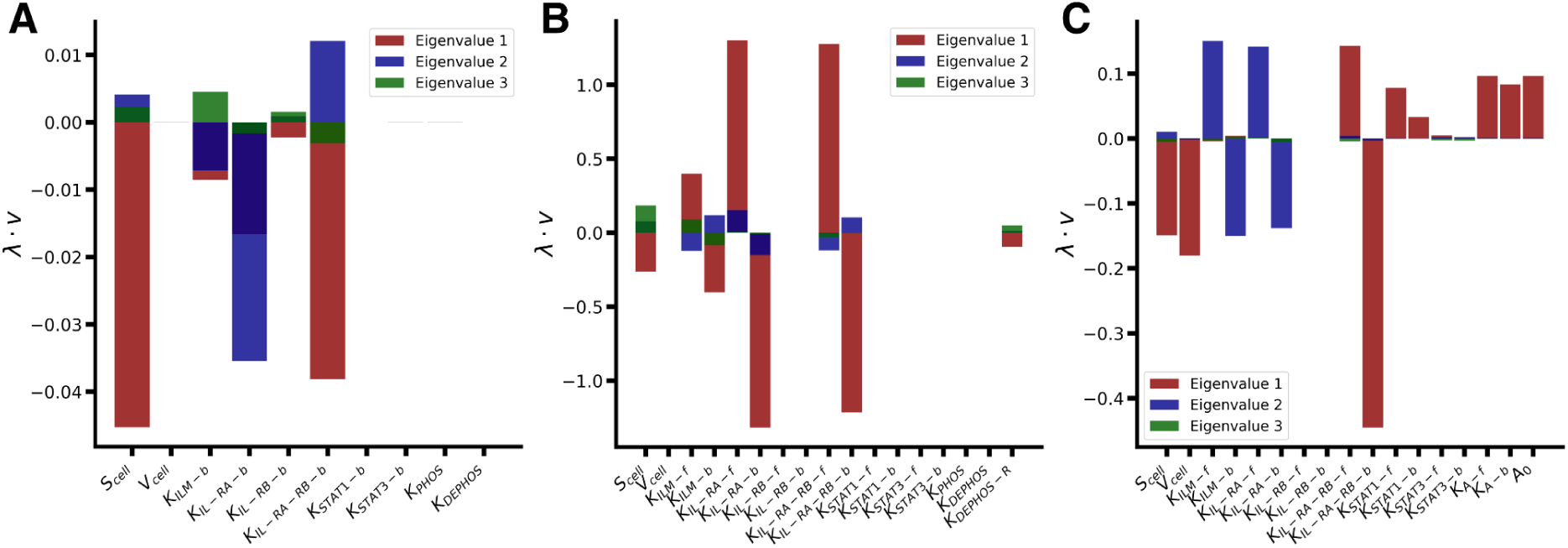
Sloppiness analysis on the 3 proposed models. **(A-C)** Eigenvalue-weighted directions of the top 3 eigenvectors with higher eigenvalues of the Laplacian matrix of the model’s sloppiness analysis for the (A) Baseline, (B) Receptor Memory, and (C) Kinetic Proofreading models. Higher absolute values indicate a bigger effect of the parameter on the model’s performance.

**Figure EV3:**
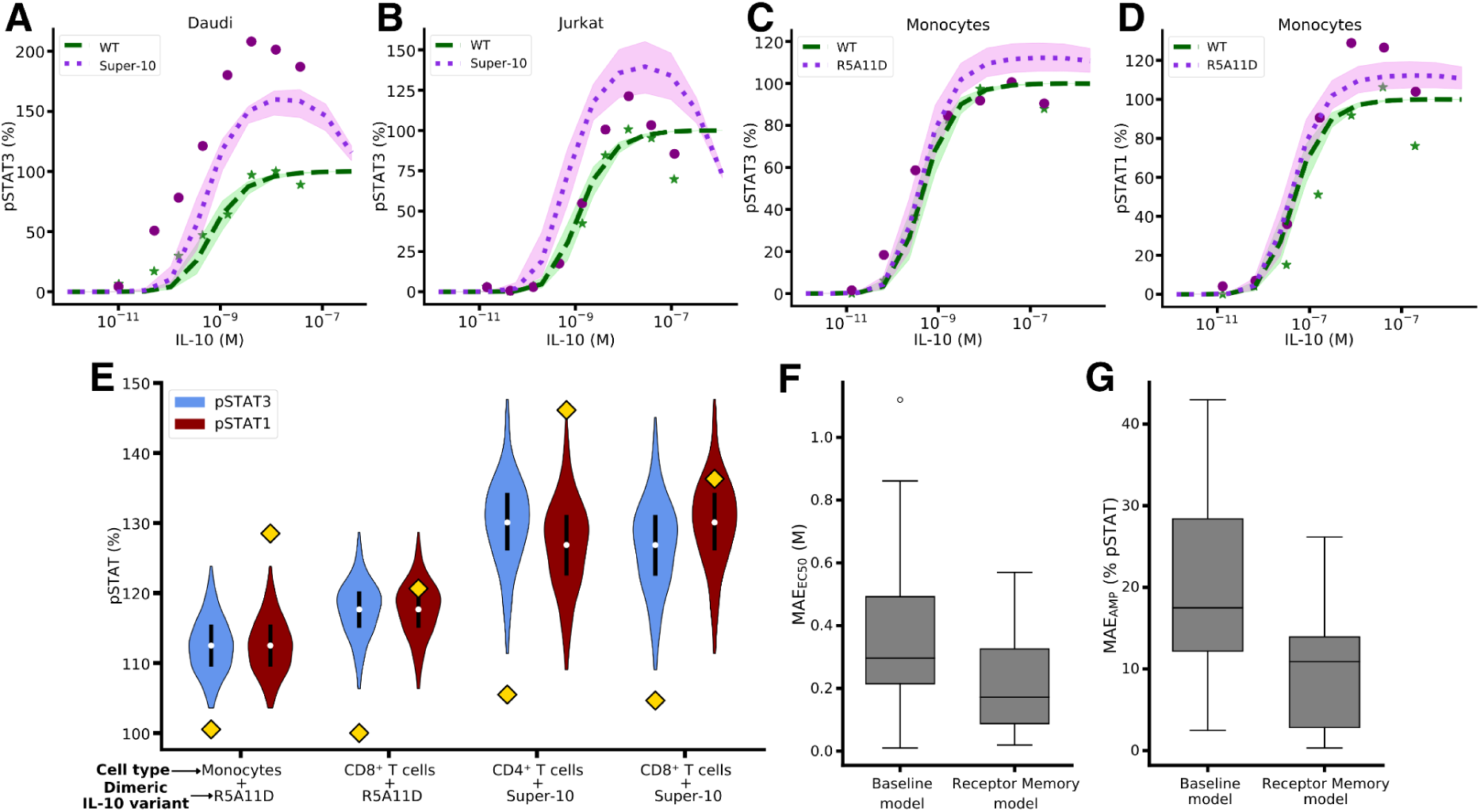
Dose-response curves on dimeric IL-10 variants for the Baseline model and model performance comparison. **(A, B)** Dose-response curves of WT IL-10 and Super-10 in the (A) Daudi and (B) Jurkat cell lines for STAT3. **(C, D)** Dose-response curves of WT IL-10 and R5A11D in monocytes for (C) STAT3 and (D) STAT1. **(E)** Violin plot of the distributions of amplitudes, defined as the maximum minus minimum pSTAT1/pSTAT3 response, of the dose-response curves in the dimeric R5A11D and Super-10 IL-10 mutants, assuming uncertainty in the cell-type-specific initial conditions. Yellow squares indicate the experimental values. As can be seen, due to the Baseline model assuming the same binding/phosphorylation for STAT1 and STAT3, it has problems fitting the amplitude of the response of mutants with enhanced IL-10RB binding at the same time for STAT1 and STAT3. **(F, G)** Mean absolute error (MAE) in the (A) EC50 (N=24) and (B) amplitude (N=14) of simulated dose-response curves with the Baseline and Receptor memory models (*p*_*EC*50_ = 0. 01, *p*_*AMP*_ = 0. 009). P-values were calculated using the Scipy implementation of the t-test with unequal variances.

**Figure EV4:**
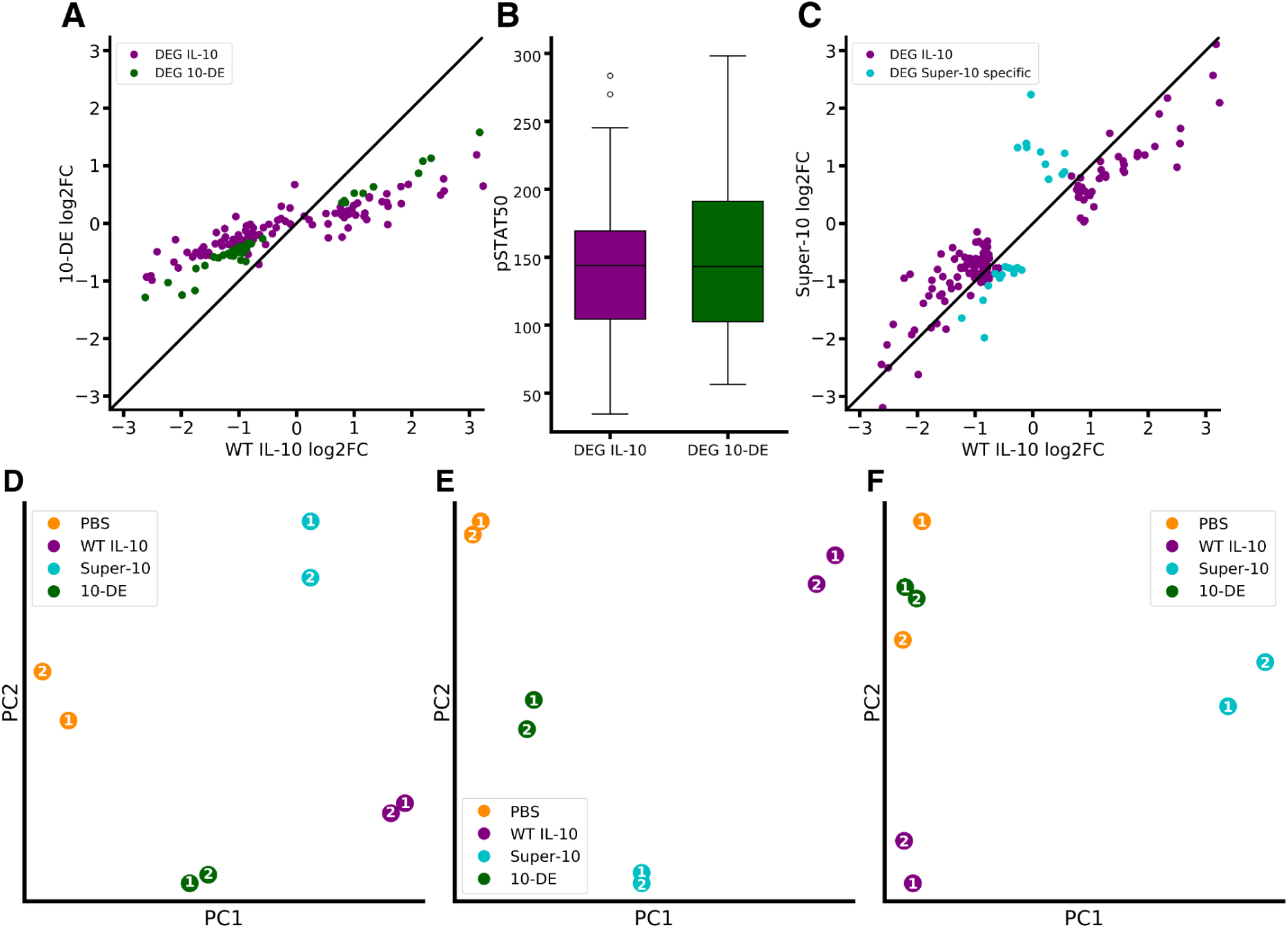
RNA response of IL-10 variants with increased or decreased STAT phosphorylation in CD8⁺ T cells. **(A)** Fold change of IL-10 DEG (shared genes responsive to at least one of the 3 IL-10 variants) in WT IL-10 and a variant with reduced STAT phosphorylation (10-DE). The black diagonal line indicates no differential transcriptional effect between WT and 10-DE. **(B)** Boxplot of the distribution of gene pSTAT thresholds (pSTAT50) among genes shared between WT IL-10 and 10-DE responses (*N* = 21) and genes only activated/repressed in WT IL-10 (*N* = 49) (*p* = 0. 71). The p-value was calculated using the Scipy implementation of the Mann–Whitney U test. **(C)** Fold change of IL-10 DEG in WT IL-10 and a variant with increased STAT phosphorylation (Super-10). **(D,E,F)** Principal component analysis of PBS, WT IL-10, and the mutants 10-DE, Super-10 on (D) genes induced by 10-DE, (E) shared genes responsive to WT IL-10 and Super-10, and (F) genes exclusively induced by Super-10. Amount of variance explained by each of the selected components: (D) (*PC*1 = 0. 96, *PC*2 = 0. 02), (E) (*PC*1 = 0. 99, *PC*2 = 0. 01), (F) (*PC*1 = 0. 98, *PC*2 = 0. 02).

**Figure EV5:**
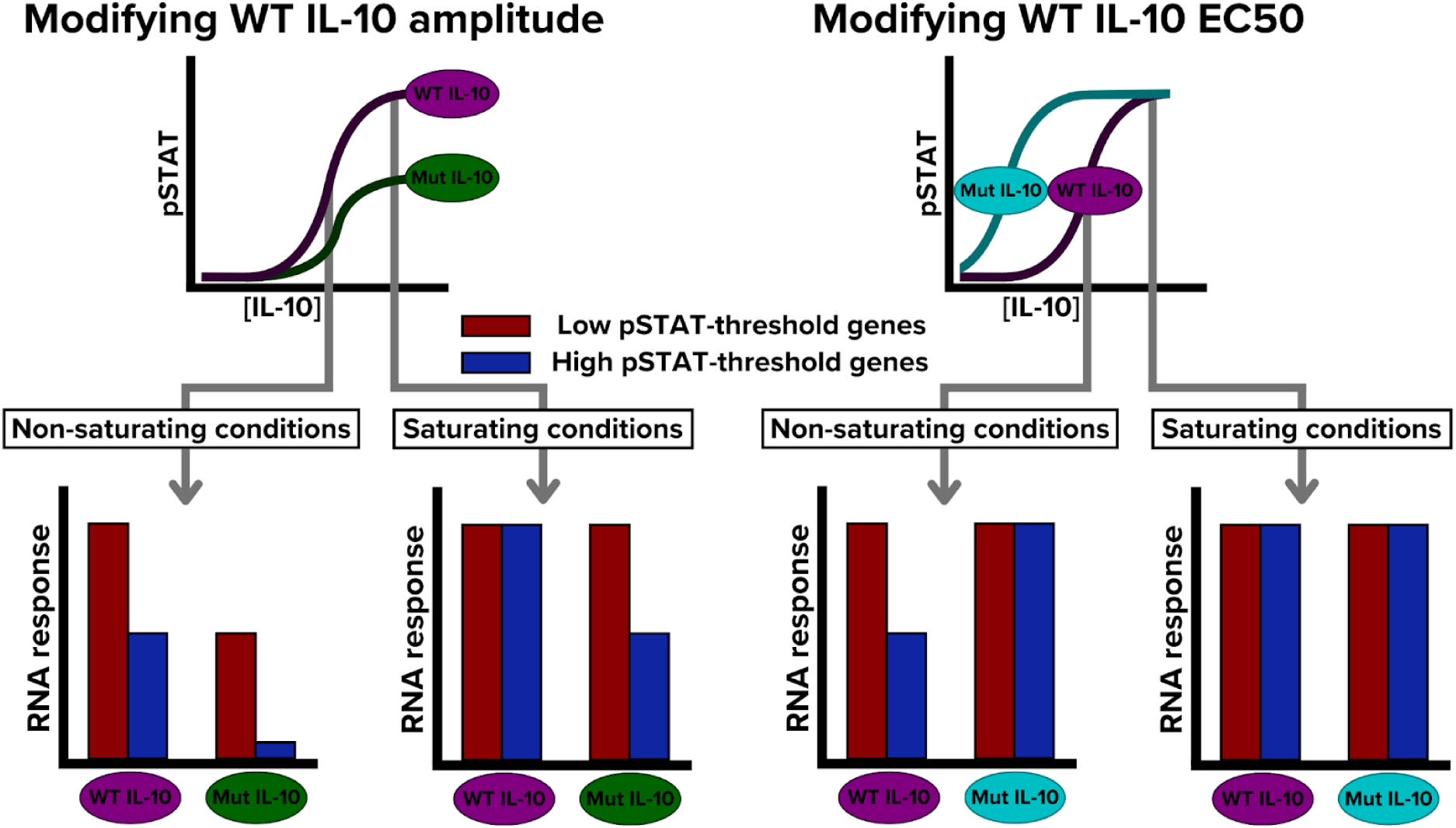
Possible effects of different IL-10 mutants on the downstream RNA response. Schematic illustrating how IL-10 mutants that alter EC50 or response amplitude of the IL-10/pSTAT curve could impact downstream RNA expression under non-saturating and saturating IL-10 concentrations. Compared to WT IL-10, mutants that modify the pSTAT amplitude could produce qualitatively distinct RNA responses at both non-saturating and saturating conditions. However, mutants that mainly alter the EC50 of the IL-10/pSTAT curve are expected to produce RNA responses similar to WT IL-10, but at different IL-10 concentrations.

## Data availability

The datasets and computer code produced in this study are available in the following databases:

- All code used in this work is available via GitHub in: https://github.com/Quim98/IL10_model_publication
- Raw proteomics data is available through the PRIDE database under the accession number: PXD076154
- Processed data needed for simulating the model and model predictions are available via Zenodo under the DOI: 10.5281/zenodo.19220929

## Author contributions

Conceptualization: QM-B, JG-O, LS; Methodology: QM-B, CS-M, JG-O, LS; Data Curation: QM-B; Investigation: QM-B; Formal analysis: QM-B, JG-O, LS; Visualization: QM-B; Software: QM-B; Experimental assays: CS-M; Validation: QM-B, JG-O, LS; Writing - original draft: QM-B; Writing - review & editing: QM-B, CS-M, JG-O, LS; Supervision: JG-O, LS; Funding acquisition: QM-B, LS; Project administration: QM-B; Resources: LS. All authors read and approved the final manuscript.

## Acknowledgements

We thank Miquel Anglada-Girotto and Rahma Hamdani for the insightful discussions. The mass spectrometry analyses were performed in the CRG/UPF Proteomics Unit, which is part of the Spanish National Infrastructure for Omics Technologies (ICTS OmicsTech). We thank Nikolaos Louros from the Switch lab of VIB-KU Leuven for his work on calculating the affinity of IL-10 to IL-10RB. We appreciate all the feedback from the members of the LS and JG-O laboratories. We acknowledge support of the Spanish Ministry of Science and Innovation through the Centro de Excelencia Severo Ochoa (CEX2020-001049-S, MCIN/AEI/10.13039/501100011033), and the Generalitat de Catalunya through the CERCA programme, and to the EMBL partnership. This work was supported by funding from “Ayudas para contratos predoctorales para la formación de doctores/as 2022” (FPI: PRE2022-101948 fundded by MICIU/AEI /10.13039/501100011033 ans ESF+). This project has also received funding from the European Research Council (ERC) under the European Union’s Horizon 2020 research and innovation programme (grant agreement No 101020135; Lung Biorepair), and Plan Estatal de Investigación Científica y Técnica y de Innovación (PID2021-122341NB-I00 project funded MICIU / AEI / 10.13039 / 501100011033 / FEDER, UE). JG-O was financially supported by the Spanish Ministry of Science and Innovation, the Spanish State Research Agency and FEDER (Project Reference No. PID2024-160263NB-I00), and by the European Research Council (ERC Synergy CeLEARN-101167121).

